# MALAT1 mediates miR-206/CDC42/PAK1/Paxillin signaling axis to alleviate erectile dysfunction in rats

**DOI:** 10.1101/2024.03.21.586086

**Authors:** Longhua Luo, Zixin Wang, Xuxian Tong, Tenxian Xiong, Minggen Chen, Xiang Liu, Cong Peng, Xiang Sun

## Abstract

Erectile dysfunction (ED) is a male sexual dysfunction with a gradually increasing prevalence, and current treatments are ineffective. This study aims to find a safer and more effective treatment for erectile dysfunction. Bone marrow-derived mesenchymal stem cells (BM-MSCs) treated with VEGFA. The expression of endothelial markers vWF, VE-cadherin and eNOS were determined by overexpressing MALAT1 and CDC42, respectively. The results showed that interference with CDC42 and MALAT1 expression inhibited the differentiation of BM-MSCs to ECs and the expression of endothelial marker-related proteins. Moreover, MALAT1 induced differentiation of BM-MCs to ECs through the CDC42/PAK1/Paxillin pathway was explored by transfecting si-MALAT1 and overexpressing CDC42 as well as PAK1 in the BM-MSCs. Next, miR-206 was overexpressed within BM-MCs to determine CDC42 expression. The binding sites of MALAT1, miR-206 and CDC42 were reported using luciferase. it was found that the MALAT1 competes with CDC42 3’-UTR for binding miR-206, which in turn is involved in differentiating BM-MSCs to ECs. Finally, DMED rat modeling success was demonstrated by APO experiments. MALAT1 overexpression-modified BM-MSCs had significant therapeutic effects in DMED rats.

## Introduction

Erectile dysfunction induced by diabetes is called diabetes mellitus erectile dysfunction (DMED).^1^ The incidence of DMED is increasing year by year, and it has become an important factor affecting men’s sexual function. However, the underlying mechanisms remain poorly understood. The mechanisms of ED complication in diabetic patients are complex. Sildenafil, tadalafil, and phosphodiesterase type 5 (PDE5) inhibitors are commonly used to prevent and control the disease in western medicine, but only improve local symptoms.^2,3^ Moreover, DMED patients are found to be resistant to PDE5 inhibitors.^4^ However, these drugs were limited use because the adverse effects. Therefore, finding safer and more effective ED treatments is especially urgent.

Bone marrow-derived mesenchymal stem cells (BM-MSCs) have been widely used in research for the treatment of neurological injuries because of their fast amplification, multidirectional differentiation potential, high regenerative capacity, low rejection, and the ability to induce differentiation. Our previous study confirmed that transplantation of BM-MSCs could effectively improve DMED, and the therapeutic effect of BM-MSCs was further improved by controlling the expression levels of some genes in BM-MSCs.^5–7^ Meanwhile, previous studies showed that overexpression of MALAT1 in BM-MSCs facilitated differentiation toward endothelial cells (ECs), whereas interference with MALAT1 inhibited differentiation of BM-MSCs toward ECs. Therefore, we will deeply investigate the mechanism of action of MALAT1 in regulating the differentiation of BM-MSCs toward ECs.^7^

The Rho GTPase is an important member of the Ras protein superfamily and is involved in the regulation of a variety of cellular events. Rho, Rac and Cell division cycle 42 (CDC42) are the most studied Rho GTPases.^8^ The study showed that up-regulation of CDC42 expression and increase of its activity contribute to angiogenesis in cultured endothelial cells *in vitro.*^9^ In addition, inducing elevated levels and activation of CDC42 contribute BM-MSCs to migrate to the site of skin injury and promoting wound healing.^10^ However, whether CDC42 contributes to the differentiation of BM-MSCs to endothelial cells has not been reported.

Paxillin is an important cell adhesion factor involved in a variety of cellular events such as cell proliferation, adhesion, and migration. It is a downstream target of p21 activated kinase 1 (PAK1), a conserved class of serine/threonine kinases.^11^ It is capable of inducing phosphorylation of Paxillin at several sites such as serine 273 and serine 258.^12,13^ Initially, PAK1 was recognized as the major effector protein of CDC42 and Rac1 in the Rho GTPase family, and it is able to participate in endothelial cell proliferation, migration, and angiogenesis by binding to CDC42 and Rac1 to form the Rac1/CDC42/PAK1 complex.^14–16^ Studies have shown that in VEGFA-treated human venous endothelial cells, the expression level of Paxillin and its phosphorylation level increased with the increase of VEGFA dose, and the expression level of CDC42 also increased.^17,18^ Meanwhile, several studies have indicated that Paxillin is involved in the regulation of endothelial cell proliferation, migration and angiogenesis.^19^ Our previous study pointed out that MALAT1 promotes the differentiation of BM-MSCs to endothelium by upregulating VEGFA expression. Therefore, we hypothesized that the CDC42/PAK1/Paxillin signaling pathway was activated in VEGFA-induced BM-MSCs.

With the development of exogenous gene transfection technology, it has become possible to transplant stem cells into tissues as gene therapy vectors to treat DMED-related diseases. In this study, the effect of implanting BM-MSCs modified by MALAT1 into penile tissues of DMED rats for the treatment of ED was studied, and the possible mechanism behind their efficacy was examined.

## Results

### Establishment of a DMED rat model and changes in endothelial marker-related protein and gene expression in cavernous tissue

There were no significant differences in baseline body weight and fasting glucose between groups before modeling. After injection of STZ, fasting glucose increased significantly and body weight was significantly lower than that of the control group (Figure 1 A-B). The number of erections in DMED rats induced by APO was significantly lower than control group (Figure 1 C). The ICP/MAP ratio was decreased in DMED rats compared with the control group (Figure 1 D). Changes in the expression of endothelial markers vWF, VE-cadherin, and eNOS-related genes in the spongiotic tissues of rats were detected by RT-qPCR and western blot. The results revealed that the expression levels of vWF, VE-cadherin, and eNOS were significantly down-regulated in DMED rats compared with control rats (Figure 1 E-F). The results showed that DMED rats modeled successfully.

**Figure 1.**
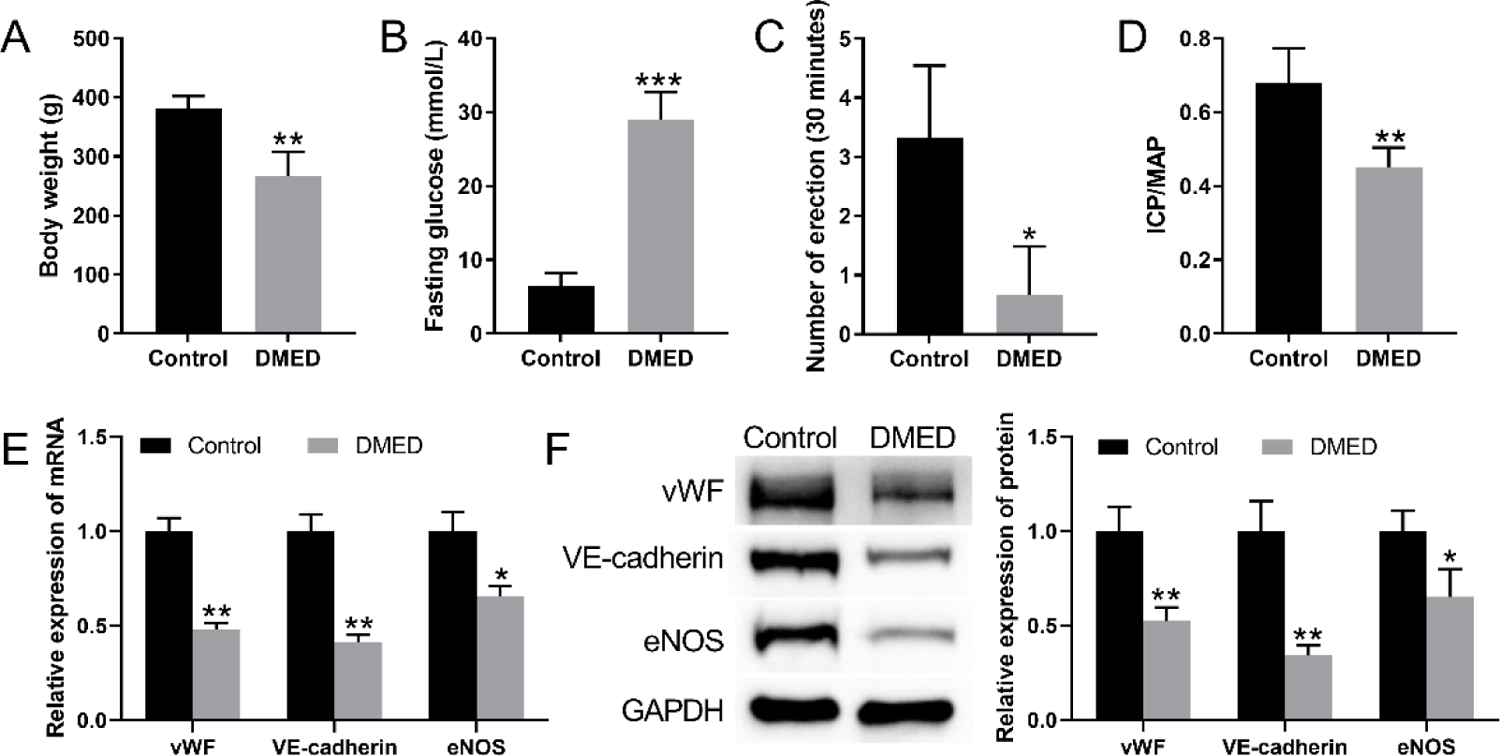
Establishment of a DMED rat model and changes in endothelial marker-related protein and gene expression in cavernous tissue. (A) Body weight of rats. (B) Fasting blood glucose of rats. (C) APO experiment to assess erectile function rats. (D) ICP/MAP ratio of rats. (E) The expression of endothelial markers vWF, VE-cadherin, and eNOS-related genes in spongiotic tissues of rats were detected by RT-qPCR. (F) The expression of endothelial markers vWF, VE-cadherin, and eNOS-related genes in spongiotic tissues of rats were detected by western blotting. \**P*<0.05, \*\**P*<0.01, \*\**P*<0.001 vs Control.

### Exploration of the functions of MALAT1 and CDC42 in VEGFA-induced differentiation of BM-MSCs to ECs

After different doses of VEGFA induced the differentiation of BM-MSCs to ECs, the expression levels of cellular endothelial markers vWF, VE-cadherin, and eNOS were found to be gradually increased with the incremental of VEGFA dose by western blotting (Figure 2 A). RT-qPCR detection revealed that the expression levels of the MALAT1 gene were gradually up-regulated with the incremental of VEGFA dose (Figure 2 B). Based on the above experiments, we chose to induce BM-MSCs with 50 ng/ml of VEGFA and transfected si-NC and si-MALAT1 in the cells, respectively. The results of RT-qPCR showed that the expression of MALAT1 was significantly reduced in the transfected si-MALAT1 group compared with the transfected si-NC group (Figure 2 C). As shown in Figure 2 D, the expression of vWF, VE-cadherin, and eNOS were up-regulated with the transfected si-MALAT1. Migration assays were performed by transwell chambers. In representative migratory images, we found that BM-MSCs showed a prominent migratory response toward VEGFA. However, transfection si-MALAT1 was induced migration (Figure 2 E).

**Figure 2.**
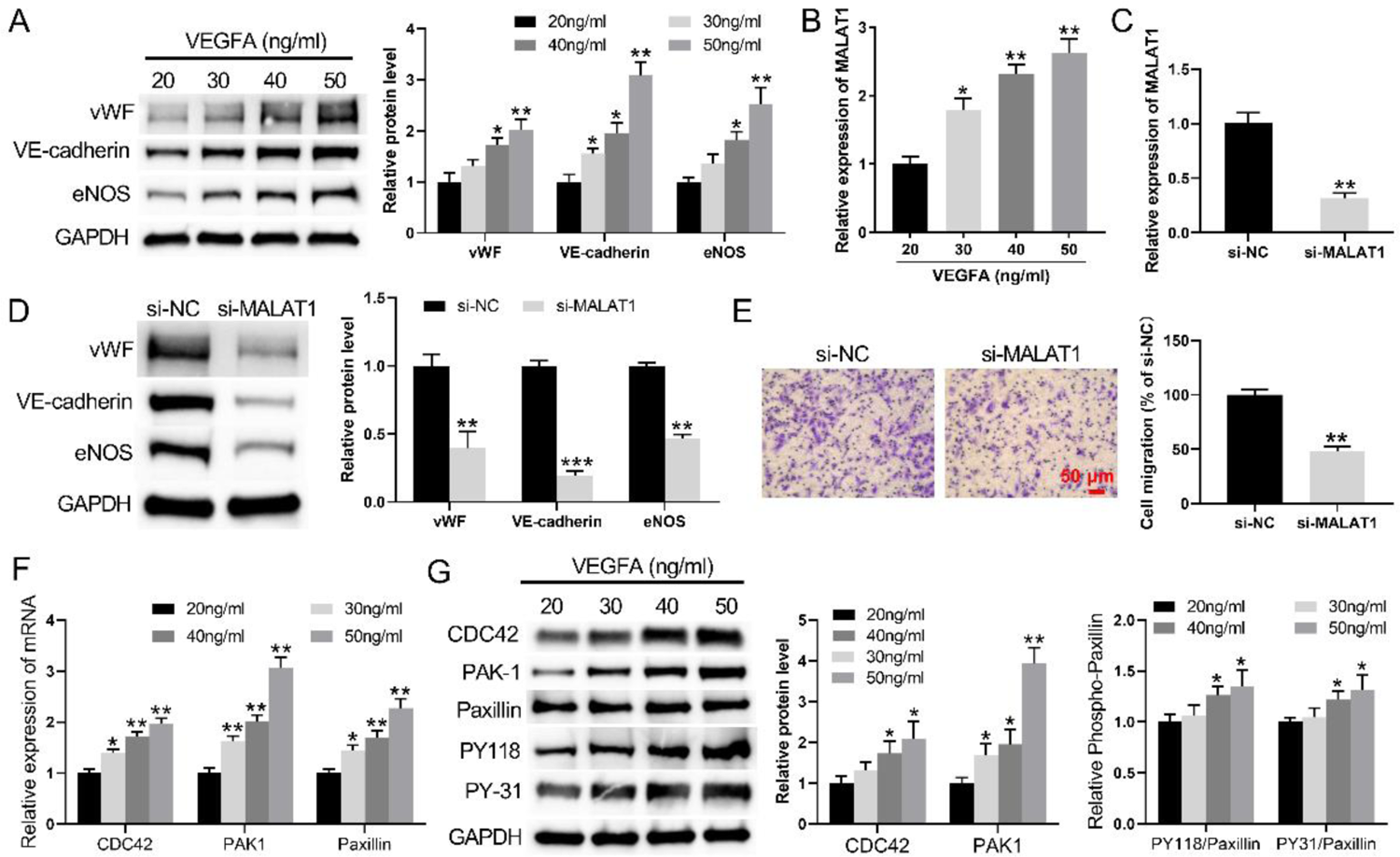
Functional exploration of MALAT1 in VEGFA-induced differentiation of BM-MSCs to endothelial cells. (A) Western blot was used to detect the levels of vWF, VE-cadherin, and eNOS protein expression in BM-MSCs induced to differentiate into endothelial cells by different doses of VEGFA treatment for 7 days. (B) RT-qPCR was used to detected the levels of MALAT1 gene expression in BM-MSCs induced to differentiate into endothelial cells by different doses of VEGFA treatment for 7 days. (C) RT-qPCR was used to detect changes in MALAT1 gene expression in BM-MSCs transfected with si-MALAT1. (D) Western blot was used to detect changes in vWF, VE-cadherin, and eNOS protein expression after transfection of si-MALAT1 within VEGFA-treated BM-MSCs. (E) Transwell assay to detect the migratory capacity of BM-MSCs. (F) RT-qPCR was used to detected the levels of CDC42, PAK1 and Paxillin gene expression in BM-MSCs induced to differentiate into endothelial cells by different doses of VEGFA treatment for 7 days. (G) Western blot was used to detect the levels of CDC42, PAK1, Paxillin and paxillin phosphorylation protein expression in BM-MSCs induced to differentiate into endothelial cells by different doses of VEGFA treatment for 7 days. \**P*<0.05, \*\**P*<0.01, \*\*\**P*<0.001 20 ng/mL group or si-NC.

Additionally, RT-qPCR and western blotting were examined the impact of different doses of VEGFA on CDC42, PAK1, Paxillin and Paxillin phosphorylation (PY118 and PY-31) expression levels in BM-MSCs. After different doses of VEGFA induced the differentiation of BM-MSCs to ECs. It was found that the expression levels of cellular endothelial markers CDC42, PAK1 and Paxillin were found to be gradually increased with the incremental of VEGFA dose by RT-qPCR (Figure 2 F). Moreover, the expression levels of cellular endothelial markers CDC42, PAK1, Paxillin, PY118 and PY-31 were found to be gradually increased with the incremental of VEGFA dose by western blotting (Figure 2 G). Then, interference with CDC42 expression within 50ng VEGFA-induced BM-MSCs. RT-qPCR and Western blotting assays revealed that the expression level of CDC42 was significantly reduced in the transfected si-CDC42 group compared with the transfected si-NC group (Figure 3 A-B). Transwell assay results showed that compared with cells transfected with si-NC, cell migration was significantly reduced after transfection with si-CDC42 in cells after VEGFA treatment of BM-MSCs (Figure 3 C). Finally, compared with cells transfected with si-NC, the expression of vWF, VE-cadherin and eNOS levels down-regulated with the transfected si-CDC42 (Figure 3 D). All these data indicated that interference with CDC42 and MALAT1 expression inhibited the differentiation of BM-MSCs to ECs and the expression of endothelial marker-related proteins.

**Figure 3.**
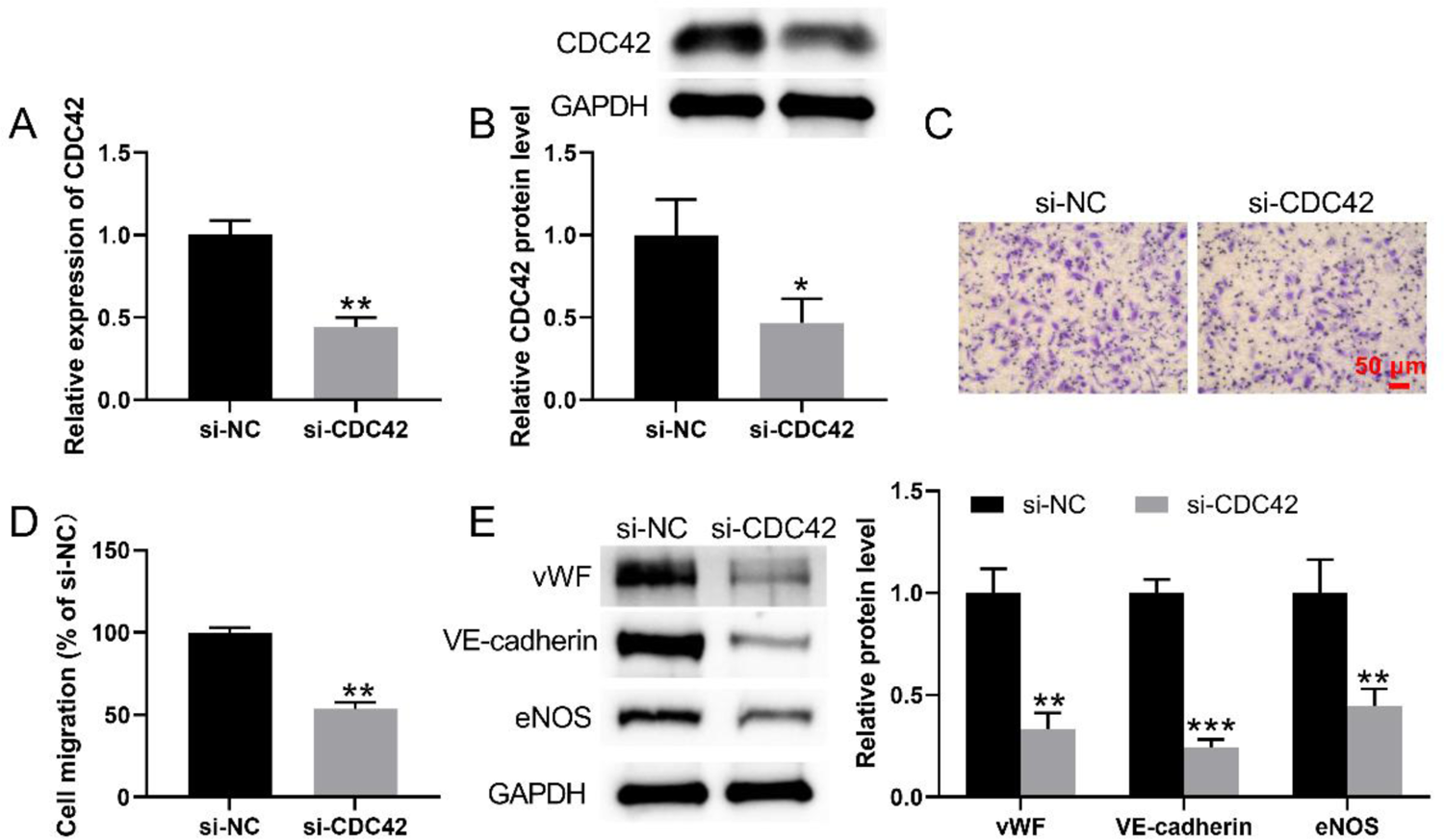
Functional exploration of CDC42 in VEGFA-induced differentiation of BM-MSCs to endothelial cells. (A) RT-qPCR was performed to detect CDC42 gene expression level. (B) Detection of CDC42 protein expression levels was detected by western blot. (C) Transwell assay to detect the migratory capacity of BM-MSCs. (D) The expression levels of cellular endothelial markers vWF, VE-cadherin, and eNOS were detected by Western blot. \**P*<0.05, \*\**P*<0.01 vs si-NC group.

### MALAT1 under VEGFA-treated conditions regulates the differentiation of BM-MSCs into ECs by modulating the CDC42/PAK1/Paxillin axis

Analysis of whether MALAT1 regulates the differentiation of BM-MSCs to endothelial cells through the CDC42/PAK1/Paxillin signaling pathway. We transfected si-MALAT1 within induced BM-MSCs with 50 ng/ml VEGFA. The expression of CDC42, PAK1, Paxillin and PY118 and PY-31 in the phosphorylation level of Paxillin in BM-MSCs was found to be significantly reduced after transfection with si-MALAT1 as detected by western blotting (Figure 4 A). Then, we inhibited MALAT1 while overexpressing CDC42 within 50 ng/ml VEGFA-induced BM-MSCs. The expression levels of PAK1, Paxillin, Paxillin phosphorylation and the endothelial cell markers vWF, VE-cadherin and eNOS were significantly down-regulated in the transfected si-MALAT1 group compared with the transfected si-NC group. In contrast, the expression levels of PAK1, Paxillin, Paxillin phosphorylation and endothelial cell markers vWF, VE-cadherin, eNOS were significantly up-regulated in the group with simultaneous overexpression of CDC42 by inhibition of MALAT1 within BM-MSCs compared with the group transfected with si-MALAT1 (Figure 4 B-C). Finally, we used VEGFA-induced BM-MSCs to intra-disrupt MALAT1 while overexpressing PAK1. As detected by western blot, the expression levels of PAK1, Paxillin, Paxillin phosphorylation and endothelial cell markers vWF, VE-cadherin, eNOS were significantly reduced in BM-MSCs transfected with si-MALAT1 compared with transfected with si-NC group. And transfection of si-MALAT1 while overexpressing PAK1 group reversed this phenomenon (Figure 4 D). The binding capacity between CDC42, PAK1 and Paxillin in VEGFA-treated and non-VEGFA-treated BM-MSCs was examined by protein immunoprecipitation. CDC42 binds to PAK1. PAK1 can bind to paxillin. And CDC42 in the added VEGFA group has a stronger ability to bind to PAK1. Similarity, PAK1 in the added VEGFA group has a stronger ability to bind to paxillin (Figure 4 E). Therefore, MALAT1 under VEGFA-treated conditions regulates the differentiation of BM-MSCs into ECs by modulating the CDC42/PAK1/Paxillin axis.

**Figure 4.**
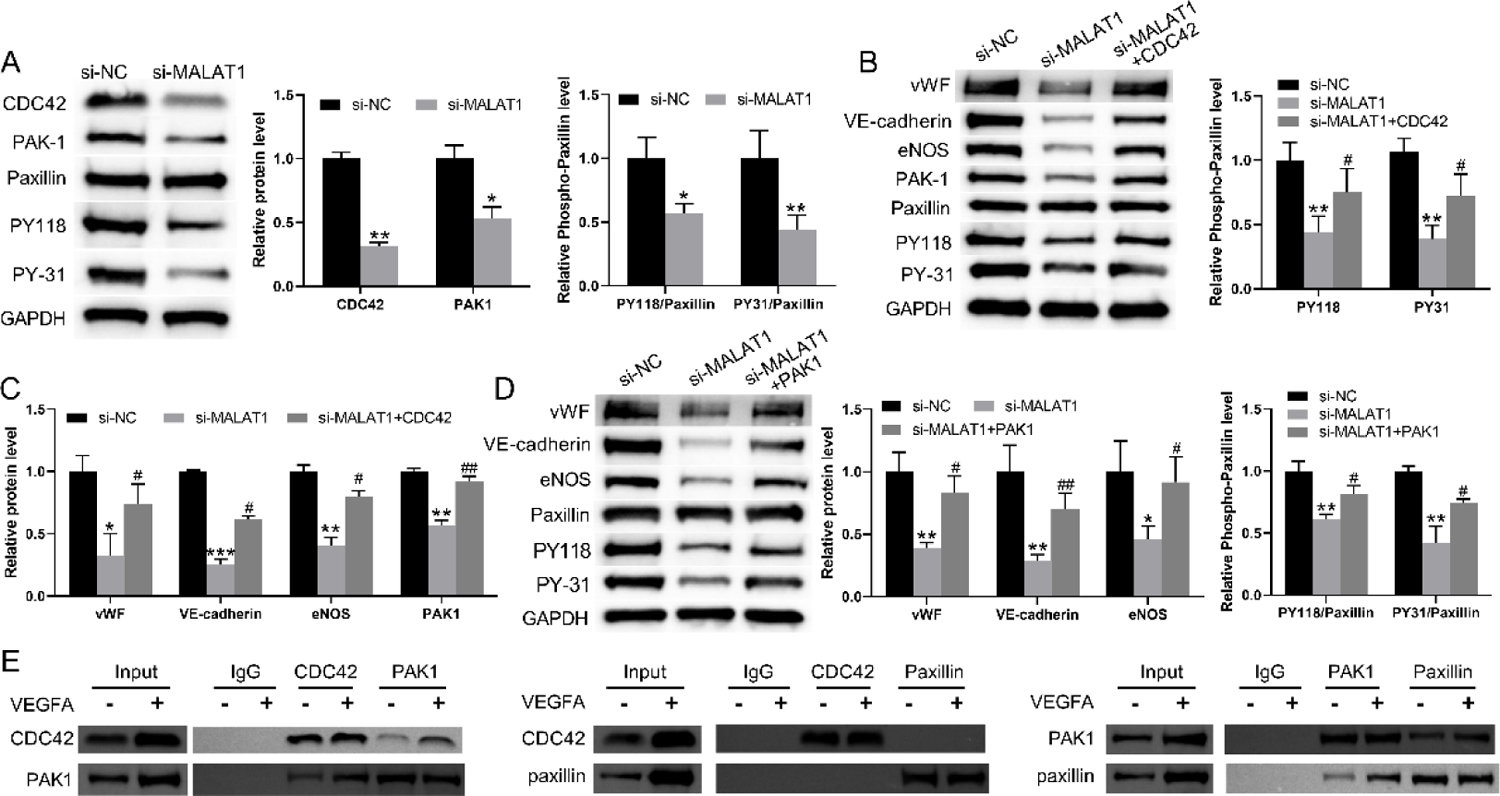
Under VEGFA-treated conditions, MALAT1 regulates the differentiation of BM-MSCs into ECs by modulating the CDC42/PAK1/Paxillin axis. (A) Western blot was used to detect the levels of CDC42, PAK1, Paxillin and Paxillin phosphorylation in BM-MSCs. (B-C) Western blot was used to detect the expression levels of PAK1, Paxillin, Paxillin phosphorylation and endothelial cell markers vWF, VE-cadherin and eNOS in BM-MSCs. (D) Western blot was used to detect the expression levels of PAK1, Paxillin, Paxillin phosphorylation and endothelial cell markers vWF, VE-cadherin and eNOS in BM-MSCs. (E) Protein immunoprecipitation was used to detect the binding capacity between CDC42, PAK1 and Paxillin in VEGFA-treated and non-VEGFA-treated BM-MSCs. \**P*<0.05, \*\**P*<0.01, \*\*\**P*<0.001 vs si-NC group, ^#^*P*<0.05, ^##^*P*<0.05 VS si-MALAT1 group.

### Mechanism of MALAT1 in regulating the differentiation of BM-MSCs to ECs

BM-MSCs were treated with VEGFA (20, 30, 40, 50 ng/ml) to induce differentiation of BM-MSCs to ECs in vitro. After RT-qPCR to detect miR-206 expression in BM-MSCs, it was found that miR-206 expression in BM-MSCs was lowest at 50 ng/ml VEGFA (Figure 5 A). Next, we overexpressed miR-206 within BM-MSCs induced by 50 ng/ml VEGFA. Detection of miR-206 transfection efficiency within BM-MSCs using RT-qPCR revealed that the expression level of miR-206 increased significantly in the overexpression miR-206 group (Figure 5 B). The western blotting results showed that the expression PAK1, Paxillin, Paxillin phosphorylation and endothelial cell markers vWF, VE-cadherin, and eNOS were decreased in BM-MSCs with transfected miR-206 mimic (Figure 5 C). To verify the interaction between MALAT1 and miR-206, CDC423’UTR and miR-206, luciferase reporter assay and RNA immunoprecipitation were carried out. The luciferase activity was notably decreased in BM-MSCs in the presence of miR-206 mimic and MALAT1 WT-1, MALAT1 WT-2 (Figure 5 D-E). Moreover, the luciferase activity was reduced in BM-MSCs in the presence of miR-206 mimic and CDC42 WT (Figure 5 F). It indicated that MALAT1 and CDC42 3’UTR interacted with miR-206. Finally, 50ng/ml VEGFA-induced intracellular overexpression of MALAT1 in BM-MSCs while overexpressing miR-206.

**Figure 5.**
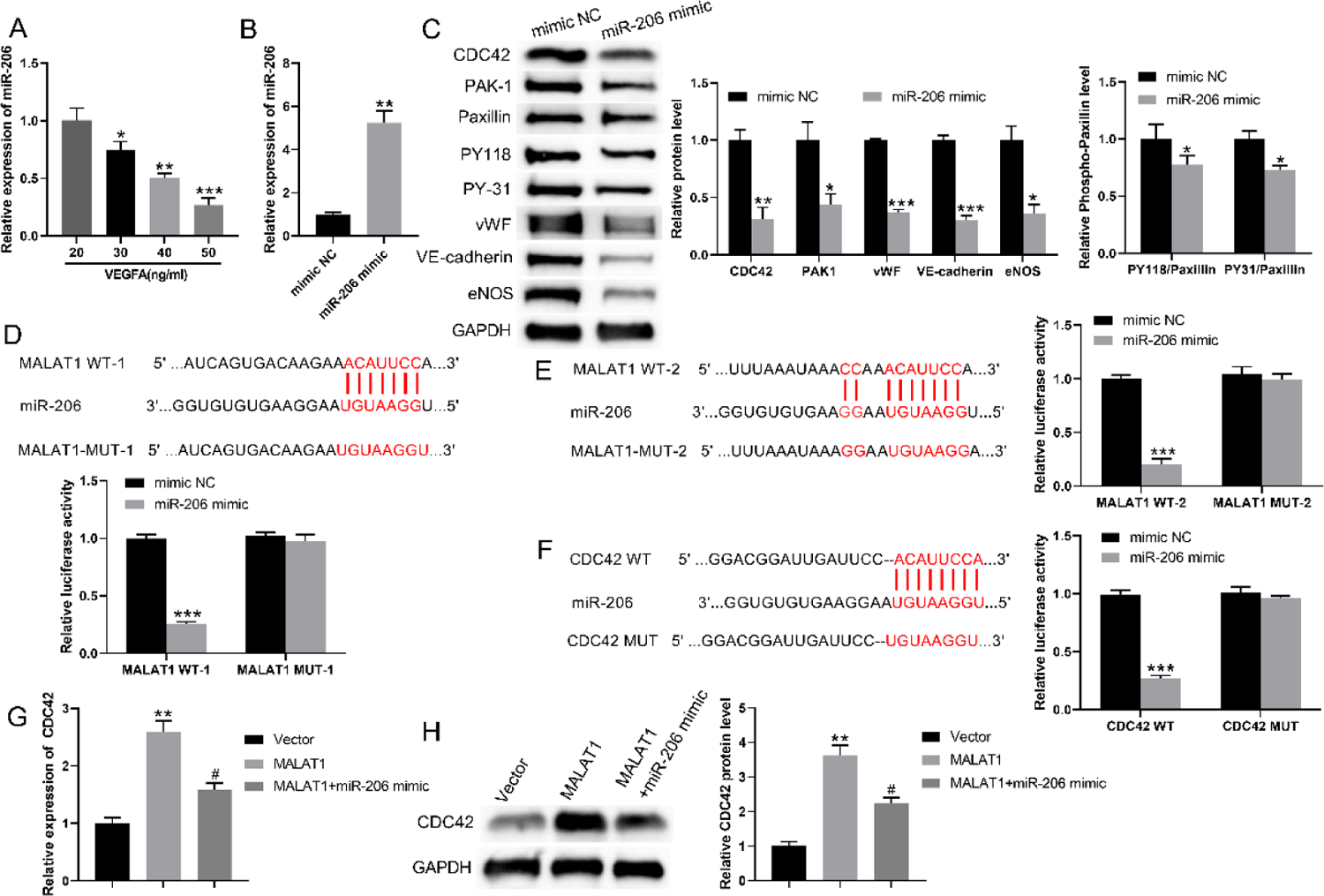
Mechanism of action of MALAT1 in regulating the differentiation of BM-MSCs to endothelial cells. (A) RT-qPCR detection of miR-206 expression changes within BM-MSCs. (B) RT-qPCR was performed to detect miR-206 expression level. (C) Western blot was used to detect the protein expression levels of PAK1, Paxillin, Paxillin phosphorylation and endothelial cell markers vWF, VE-cadherin and eNOS. (D-E) Schematic representation of the two binding sites of MALAT1 and miR-206 and luciferase assay to detect the binding between the two binding sites of MALAT1 and miR-206. (F) Schematic diagram of CDC423’UTR and miR-206 binding site and luciferase assay to detect the binding between CDC423’UTR and miR-206 binding site. (G) Detection of CDC42 gene levels within BM-MSCs was performed by RT-qPCR. (H) Detection of CDC42 protein expression levels within BM-MSCs by Western blot. \**P*<0.05, \*\**P*<0.01, \*\*\**P*<0.001 vs 20 ng/mL VEGFA group or mimic NC/Vector. ^#^*P*<0.05 vs MALAT1 group.

The expression level of CDC42 was significantly increased in the MALAT1 overexpression group compared with the overexpression control group. And the expression level of CDC42 was significantly decreased in the MALAT1 and miR-206 simultaneous overexpression group compared with the MALAT1 overexpression group. But not lower than the CDC42 expression level in the overexpression control group (Figure 5 G-H). All these data indicated that MALAT1 competes with CDC42 3’-UTR for binding miR-206, which in turn is involved in the differentiation of BM-MSCs to ECs.

### Involvement of MALAT1 in the implantation of BM-MSCs in the repair of erectile function in DMED rats

Finally, sixty male SD rats were randomly divided into 6 treatment groups (n = 10/group): control, DMED, DMED rats implanted with PBS, BM-MSCs, BM-MSCs transfected with MALAT1, and BM-MSCs combined with sildenafil treatment, respectively. Changes in body weight and fasting blood glucose were significantly lower in rats in the remaining groups compared to the control group. However, DMED rats implanted with BM-MSCs, BM-MSCs transfected with a lentivirus overexpressing MALAT1, and BM-MSCs combined with sildenafil treatment had no significant effect on body weight and blood glucose in diabetic rats (Figure 6 A-B). Then, APO experiments were performed to assess erectile function in each group of rats. The results showed that the number of erections in DMED rats and DMED rats implanted with PBS induced by APO was significantly lower than control group. However, DMED rats implanted with BM-MSCs, BM-MSCs transfected with MALAT1, and BM-MSCs combined with sildenafil treatment were obviously increased the number of erections (Figure 6 C). In addition, the ICP/MAP ratio was decreased in DMED and DMED rats implanted with PBS groups compared with the control group. Compared with the DMED model group, the ICP/MAP ratio was significantly higher in the three treatment groups treated with BM-MSCs (Figure 6 D).

**Figure 6.**
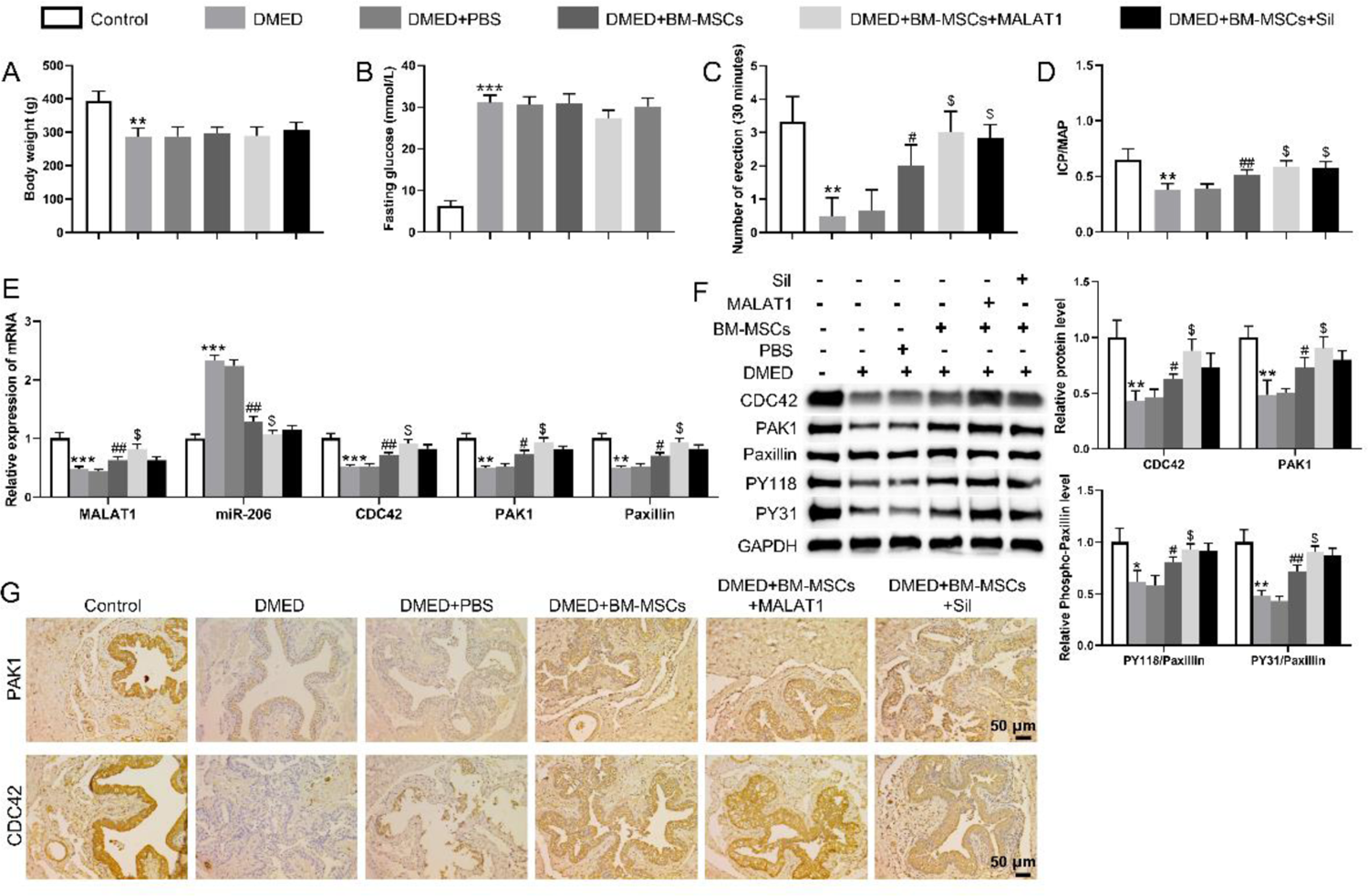
Repairing effect of MALAT1 on erectile function in DMED rats. (A) Changes in body weight of rats in each group. (B) Changes in fasting blood glucose in rats of all groups. (C) The APO assay assessed the erectile function of rats in each group. (D) ICP/MAP ratio of rats in each group. (E) RT-qPCR was used to detect the levels of MALAT1, miR-206, CDC42, PAK1, and Paxillin genes in the penile corpus cavernosum tissues of rats in each group. (F) Western blot assay was used to detect the levels of CDC42, PAK1, Paxillin and Paxillin phosphorylated proteins in the penile corpus cavernosum tissues of rats in each group. (G) Representative pictures of CDC42 and PAK1 were detected by immunohistochemistry in cavernosum tissues of rats in each group. \*\**P*<0.01, \*\*\**P*<0.001 vs control group; ^#^*P*<0.01, ^##^*P*<0.01 vs DMED+ PBS group; ^$^*P*<0.05 vs DMED+BM-MSCs.

RT-qPCR was used to detect the levels of MALAT1, miR-206, CDC42, PAK1, and Paxillin genes in the penile corpus cavernosum tissues of rats in each group. The results revealed that the expression levels of MALAT1, CDC42, PAK1, and Paxillin were significantly down-regulated in DMED rats compared with control rats. When differently treated BM-MSCs were implanted in DMED rats, it was found that the expression levels of MALAT1, CDC42, PAK1, and Paxillin were significantly up-regulated compared with DMED rats. However, the expression levels of miR-206 was increased in DMED rats. This was reversed when differently treated BM-MSCs were implanted in DMED rats (Figure 6 E). Then, the expression of CDC42, PAK1, Paxillin and Paxillin phosphorylation were detected by western blotting. The results showed that the expression of CDC42, PAK1, Paxillin, PY31, and PY118 were decreased in DMED rats compared with control group. However, when differently treated BM-MSCs were implanted in DMED rats, the expression of CDC42, PAK1, Paxillin, PY31, and PY118 were up-regulated compared with DMED rats (Figure 6 F). Finally, the expression of CDC42 and PAK1 was detected by immunohistochemistry. CDC42 and PAK1 positive cells were more expressed in the control group and mainly distributed in perivascular endothelial cells. The mean optical density values of CDC42 and PAK1 in the DMED model group were significantly lower than those in the control group. After implantation of BM-MCs with different treatments in the DMED, the mean optical densities of CDC42 and PAK1 were significantly higher compared with those in the DMED group (Figure 6 G). Among them, MALAT1 overexpression-modified BM-MSCs had significant therapeutic effects in DMED rats.

## Discussion

In recent years, with the development of social economy and the change of people’s lifestyle, high-fat diet and lack of exercise have led to more and more people suffering from DM. The increasing prevalence of DM and the younger age of the patients are more likely to induce ED, which has become an important factor affecting the health of men’s lives and the harmony of their families, and has attracted great attention from sociologists and medical researchers.^20^ Medication is a common treatment, but the therapeutic effect is not satisfactory. Mesenchymal stem cells for ED is a promising approach and many studies have been conducted.^21,22^ However, more investigations are still needed before this approach is feasible and effective.^23,24^ In this study, we found that treatment with BM-MCs overexpressing MALAT1 significantly attenuated ED in rats, suggesting that MALAT1 plays an important role in ED improvement.

MALAT1 is a lncRNA that is highly expressed in lung cancer tissues and has been reported to be associated with tumor cell proliferation and metastasis.^25^ However, the MALAT1 function in erectile function and mesenchymal stem-cell differentiation remains unclear. In a previous report from our laboratory, the expression level of MALAT1 in rat penile tissues was significantly increased after transplantation treatment of BM-MSCs. And interfering with MALAT1 expression in BM-MSCs followed by transplantation treatment significantly reduced the treatment effect of BM-MSCs. In this work, the mechanism of action of MALAT1 in regulating the differentiation of BM-MSCs to endothelial cells was deeply investigated. We found that MALAT1 up-regulates CDC42 levels by inhibiting miR-206 expression and promotes PAK1 expression and induced paxillin phosphorylation, which in turn contributes to the differentiation of BM-MSCs to endothelial cells. MicroRNAs are non-coding RNAs with a length of only 19-30 nucleotides, which can degrade mRNAs or inhibit their expression through translational repression.^26,27^ In recent years, microRNAs have been found to play a wide range of physiological regulatory roles.^28^ Among the many target genes predicted by Targetscan Human, CDC42 attracted our attention. We found that CDC42 3’-UTR has a binding site to miR-206.

CDC42 is a protein with GTPase activity in the Rho family, which is involved in various physiological processes, including cell division, intracellular transport, gene transcription, cell cycle regulation, and regulates cell growth and stability.^29–31^ We overexpressed miR-206 within VEGFA-treated BM-MSCs to detect the expression levels of CDC42 gene and protein in the cells. It was found that the expression of CDC42 was down-regulated in cells transfected with miR-206. In addition, CDC42 can activate downstream proteins, such as PAKs, ACK1, IQGAPs and P13Ks, which are closely related to cell invasion, migration, proliferation, and neovascularization.^32,33^ P21-activated kinase-1 (PAK1) is an important effector protein of CDC42 and is closely related to cell motility.^34^ Meanwhile, several studies have indicated that Paxillin is involved in the regulation of endothelial cell proliferation, migration and angiogenesis.^35,36^ Overexpression of CDC42 in BM-MSCs cells detected the expression of intracellular PAK1. We found that the expression of PAK1 was significantly elevated in cells overexpressing CDC42 compared to cells transfected with si-MALAT1. Subsequently, PAK1 was overexpressed in BM-MSCs cells and the expression of paxillin, paxillin phosphorylation was detected. The results showed that the expression levels of PY118 and PY31 were significantly up-regulated in cells overexpressing PAK1 compared to cells transfected with si-MALAT1. As expected, our results in this study showed that overexpression of MALAT1-modified BM-MSCs effectively improves DMED rats. MALAT1 upregulates CDC42 levels by inhibiting miR-206 expression, promotes PAK1 expression and its induced phosphorylation of paxillin, and induces upregulation of the expression levels of endothelial markers vWF, VE-cadherin, and eNOS. This in turn contributes to the differentiation of BM-MSCs to ECs and improves DMED in rats.

## Conclusion

Collectively, overexpression of MALAT1-modified BM-MSCs effectively ameliorated DM-induced erectile dysfunction in rats. MALAT1 up-regulates CDC42 levels by inhibiting miR-206 expression, promotes PAK1 expression and its induced phosphorylation of paxillin, which then contributes to the differentiation of BM-MSCs into ECs and ameliorates DMED in rats.

## Materials and methods

### Animals and treatment

Sixty male SD rats aged 10 weeks were purchased from Beijing Weitong Lihua laboratory animal science and technology Co., Ltd. And the mating test confirmed that they had normal erectile function. The rats were randomly divided into experimental and control groups after 1 week of free-feeding and water-acclimatization. Rats were injected intraperitoneally with 1% streptozotocin (Solarbio, Beijing, China) to construct a DM rat model, and blood glucose levels were measured after 72 h. Rats with blood glucose levels higher than 16.7 mmol/L were considered successful DM models. Diabetic rats were fed for 8 weeks to develop ED. Erectile function was assessed using the apomorphine (APO) (Solarbio, Beijing) induced erection test. Penile erection was characterized by either penile head engorgement or penile growth, and rats without penile erection were considered DMED rats. After the DMED model was constructed in the experimental group of rats, the experiments were conducted by implanting PBS, BM-MSCs, MALAT1 overexpression in BM-MSCs, and BM-MSCs combined with sildenafil (Solarbio, Beijing) treatment. This animal experiment was approved by the Ethics Committee of the First Affiliated Hospital of Nanchang University.

### Intra cavernosa pressure (ICP) and mean arterial pressure (MAP) assessment

3% sodium pentobarbital solution was injected intraperitoneally, and after confirming the anesthesia of the rats. The rats were immobilized and the abdomen was disinfected with iodophor. Two PE50 tubes connected to a pressure sensing device were placed into the left common carotid artery and the cavernous body of the penis for pressure measurement. The cavernous nerves located in the dorsal aspect of the rat prostate were isolated using a microscopic instrument. Electrical stimulation was performed with a bipolar hook from the pelvic ganglion 3-5 mm for 60 s, resulting in different degrees of penile erection. The physiological pressure signal was transduced by a physiological signal transducer and the ICP curve was recorded.

### Cell culture and treatment

The rats were killed by cervical dislocation, and the femur was isolated and the bone marrow cavity was exposed. The bone marrow cavity was repeatedly blown with DMEM (Gibco, Grand Island, NY, USA) low sugar medium. The medium containing bone marrow was added to a centrifuge tube and centrifuged at 1,000 r/min for 10 min, and the supernatant was discarded. The obtained cells were cultured at 37 ℃ in an incubator containing 5% CO_2_. The cell growth was observed by inverted microscope. BM-MSCs were treated with 20, 30, 40, and 50 ng/ml of VEGFA to induce differentiation towards endothelial cells.

### Cell transfection

The cDNA of MALAT1 or CDC42 was cloned into pcDNA3.1 vector (Invitrogen, Thermo Fisher Scientific). An empty pcDNA3.1 vector was used as a negative control (NC). GenePharma (Shanghai, China) provided miR-206 mimic/inhibitor and the corresponding controls mimic/inhibitor NC, RNA oligonucleotides for MALAT1 knockdown (si-MALAT1), RNA oligonucleotides for CDC42 knockdown (si-CDC42). Subsequently, the BM-MSCs was transfected using Lipofectamine® 3000 (Invitrogen, USA) before chemical treatment.

### RT-qPCR

Total RNA from rat cavernosum tissue was extracted by adding Trizol (Thermo Fisher Scientific), chloroform and isopropanol according to the instructions, and the extracted total RNA was used to remove DNA using the DNA Eraser Buffer kit (TaKaRa, Beijing, China). And the obtained RNA was used for reverse transcription using the PrimeScript RT Enzyme Mix I kit (TaKaRa). After the reaction was completed, primers were added for PCR amplification. The reaction conditions were set at 95℃ for 30s of pre-transformation, 95℃ for 5s of denaturation, 55℃ for 30s of annealing, and 72℃ for 30s of reaction. The reaction was fully extended and the fluorescence was collected, and the above reaction was cycled for 40 times. Total RNA was extracted from cultured BM-MSCs using RNeasy kit (Qiagen, Hilden, Germany). Real-time RT-PCR was performed as described previously. The final results were analyzed by 2^-ΔΔCT^ method.

### Western blot

BM-MSCs in logarithmic growth phase and in good condition were washed twice with PBS (Gibco, Grand Island, NY, USA). Subsequently, RIPA (Beyotime, Shanghai, China) was added to lysed the cells, and 30 μg of total protein was extracted for sampling. The cells were incubated with primary antibody (anti-vWF 1:200, anti-VE-cadherin 1:500, anti-eNOS 1:500, anti-MALAT1 1:500, anti-CDC42 1:500, anti-PAK1 1:500, anti-Paxillin 1:500, anti-PY31 1:500, anti-PY118 1:500) overnight at 4 ℃. The bands were analyzed by incubation at room temperature for 1 h with secondary antibody (goat anti-rabbit IgG 1:3,000).

Tissue proteins of penile cavernosum were taken and lysed by adding RIPA after grinding in a low temperature homogenizer, and 60 μg of samples were taken. The samples were incubated the primary antibody (anti-vWF 1:200, anti-VE-cadherin 1:500, anti-eNOS 1:500, anti-CDC42 1:500, anti-PAK1 1:500, anti-Paxillin 1:500, anti-PY31 1:300, anti-PY118 1:500) at 4 ℃ overnight. And then incubate the secondary antibody (rabbit anti-mouse IgG 1:3 000) at room temperature. Incubate at room temperature with secondary antibody (rabbit anti-mouse IgG 1:3,000) for 1 hour and analyze the bands by ImageJ software.

### Co-immunoprecipitation (Co-IP) assay

Add 40 μL protein A + G microspheres into the extracted nuclear protein, incubate at 4℃ for 1h, centrifuge at 14000×g for 2min, and then remove the supernatant. Add 10μL CDC42/PAK1/Paxillin polyclonal antibody and incubate at 4℃ overnight. Add 40μL of protein A+G microspheres, incubate at 4℃ for 2h, then centrifuge at 50,000×g for 20s, remove the supernatant, collect the microspheres, rinse with PBS, and then add 50μL of SDS Sampling Buffer containing 5% β-mercaptoethanol. Heat to 85℃ for 10 min and centrifuge at 50,000×g for 2 min, remove the supernatant for protein electrophoresis, and then perform silver staining on the electrophoresis gel. The supernatant was taken for protein electrophoresis and the electrophoresis gel was silver-stained.

### Transwell migration assay

After trypsin digestion, the cell concentration was adjusted to 5×10^4^ cells/mL in serum-free DMEM medium. And 200 μL of cell suspension was added to the upper chamber of the transwell, and 10% FBS DMEM culture medium was added to the lower chamber. The chamber was taken out 24 h later, and the cells at the bottom of the upper chamber were wiped off with a cotton swab. 4% paraformaldehyde was used for 15 min, and the cells were washed with PBS and stained with 0.5% crystal violet for 15 min, and then the cells were washed and photographed under the microscope. Five fields of view were randomly selected for counting, and the results of three replicate experiments were analyzed statistically.

### Luciferase assay

Wild-type and mutant MALAT1 (with a mutated miR-206 binding site) were cloned into the pmirGLO dual luciferase vector (Gene Pharma) to construct a dual luciferase reporter plasmid. BM-MSCs cells were co-transfected with wild-type pmirGLO-MALAT1 (or mutant) and miR-206 mimics (or negative control) using Lipofectamine 2000. Similarly, dual luciferase reporter plasmids containing CDC42-WT and CDC42-MUT were constructed. BM-MSCs cells were co-transfected with miR-206 mimics/inhibitors or their corresponding empty vectors and luciferase reporter plasmids using Lipofectamine 2000. Forty-eight hours after transfection, luciferase activity was analyzed using the dual-luciferase reporter kit (Promega, USA).

### Immunohistochemistry

The 10% formaldehyde-fixed tissues were dehydrated by an automatic dehydrator and then embedded and sectioned. The sections were deparaffinized and immersed in 3% methanol hydrogen peroxide for 10 min, washed three times with PBS for 5 min each time. The sections were heated to boiling in citrate buffer, washed with PBS, and then immersed in blocking solution for 20 min. Rabbit anti-mouse CDC42 and PAK1 were then added and incubated overnight. Biotinylated secondary antibody was added dropwise, incubated at 37℃ for 30 min, washed with PBS for 3 times. Then the sections were stained with 3, 3’-Diaminobenzidine (DAB). The slices were sealed with clear gum and photomicrographs were taken under light microscope. The results were analyzed using Image-Pro Plus 6.0.

### Statistical analysis

Each assay was performed for 3 times. Data were analyzed by SPSS 22.0 statistical software (IBM, Armonk, NY, USA) and expressed as mean ± standard deviation. Independent sample’s *t* test and one-way ANOVA were used to detect differential expression of indicators. P<0.05 was considered as a significant difference.

## Declarations Ethics statement

All protocols were authorized by the Ethics Committee of First Affiliated Hospital of Nanchang University.

## Consent for publication

Not applicable.

## Availability of data and materials

The datasets generated and/or analysed during the current study are not publicly available, but are available from the corresponding author on reasonable request.

## Acknowledgements

The authors thank all colleagues and supports from Nanchang University.

## AUTHOR CONTRIBUTION

Conceptualization: Longhua Lou, Zixin Wang. Data curation: Longhua Lou, Zixin Wang. Formal analysis: Xuxian Tong, Tenxian Xiong, Minggen Chen. Funding acquisition: Xiang Sun. Investigation: Xiang Liu, Cong Peng. Methodology: Longhua Lou, Zixin Wang. Project administration: Longhua Lou, Zixin Wang. Resources: Xiang Sun. Supervision: Xiang Sun. Writing – original draft: Longhua Lou, Zixin Wang. Writing – review & editing: Xuxian Tong, Tenxian Xiong, Xiang Sun.

